# Capturing hidden regulation based on noise change of gene expression level from single cell RNA-seq in yeast

**DOI:** 10.1101/2021.06.29.450440

**Authors:** Thoma Itoh, Takashi Makino

## Abstract

Recent progress in high throughput single cell RNA-seq (scRNA-seq) has activated the development of data-driven inferring methods of gene regulatory networks. Most network estimations assume that perturbations produce downstream effects. However, the effects of gene perturbations are sometimes compensated by a gene with redundant functionality (functional compensation). In order to avoid functional compensation, previous studies constructed double gene deletions, but its vast nature of gene combinations was not suitable for comprehensive network estimation. We hypothesized that functional compensation may emerge as a noise change without mean change (noise-only change) due to varying physical properties and strong compensation effects. Here, we show compensated interactions, which are not detected by mean change, are captured by noise-only change quantified from scRNA-seq. We investigated whether noise-only change genes caused by a single deletion of STP1 and STP2, which have strong functional compensation, are enriched in redundantly regulated genes. As a result, noise-only change genes are enriched in their redundantly regulated genes. Furthermore, novel downstream genes detected from noise change are enriched in “transport”, which is related to known downstream genes. Herein, we suggest the noise difference comparison has the potential to be applied as a new strategy for network estimation that capture even compensated interaction.

## Introduction

Cells are controlled and maintained by proteins translated from thousands of genes. These genes interact with each other to exert their functions in the cells [1]. To understand the regulatory mechanisms of gene expression, the regulatory relationships among genes have been closely examined [2]. Gene regulatory networks (GRNs) are directed graphs showing regulatory relationships of transcription factors (TFs) and their target genes. Since the GRN plays an important role in connecting genes and phenotypes, uncovering interactions contributes to understanding diverse biological phenomena. In recent years, a vast amount of expression data has rapidly accumulated through the development of high-throughput RNA-seq, resulting in data-driven GRN estimation methods, which have drawn much attention. Most GRN estimation methods are based on the assumption that perturbations in a specific gene affects downstream genes [3]. Capturing change in the expression level due to deletion of a specific gene effectively estimates molecules downstream of the deleted gene. However, the actual GRN is more complex, and may not be estimated with this simple assumption. Hundreds of downstream genes of STP2, which is a TF, have been validated by experimental approaches. Interestingly, most were detected from the double deletion of *STP1* and *STP2* [4, 5], and did not show a change in the mean expression level when only *STP2* was deleted [6]. This phenomenon would be explained by functional compensation of a *STP2* paralog. Since the lost function, upon *STP2* deletion, was compensated by *STP1*, which is the functionally redundant gene with *STP2*, the downstream genes of STP2 are not affected by the *STP2* deletion [7], resulting in no change in expression level of STP2 downstream genes.

While functional compensation provides robustness to perturbations in the GRN, it hinders estimation of downstream genes. To identify downstream genes that are not affected by upstream perturbations due to functional compensation, we focused on a change in expression noise by gene deletion. Expression noise is the stochastic variation in gene expression quantified by variance excluding the contribution of the mean in a clonal population [8]. Since expression noise is affected by the transcription or translational efficiency characterized by the physical property of a sequence [9], even functionally redundant gene pairs may show different expression noise. Furthermore, since it has been reported that expression noise propagates from upstream genes, the expression noise altered by compensation of redundant genes is expected to propagate to downstream genes and alter the expression noise of downstream genes [9]. Based on these previous reports, we hypothesize that noise change can be observed downstream of functionally compensated genes due to the physical change in upstream genes (Fig. 1).

**Figure 1.**
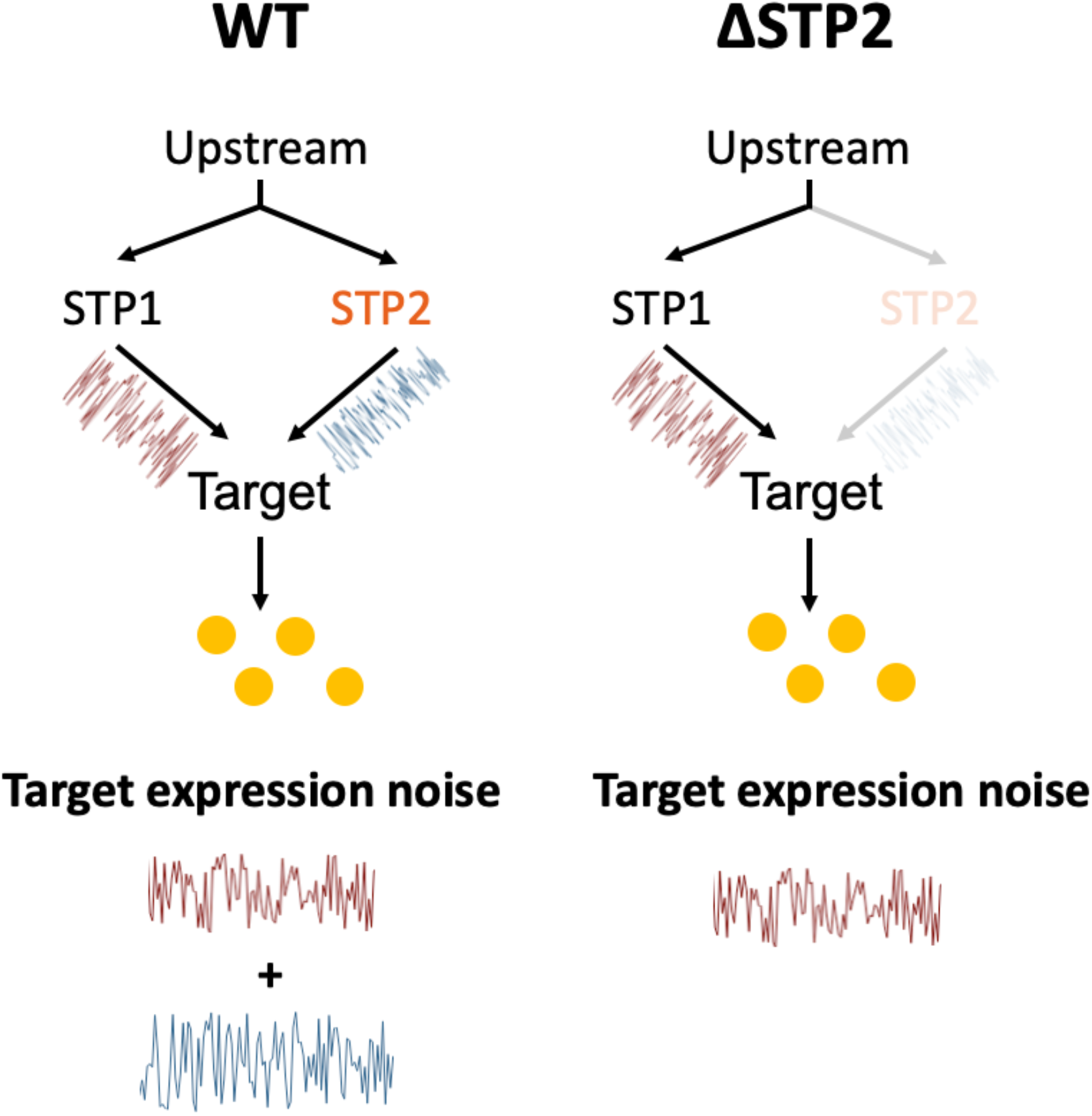
Noise difference detects functional compensation. STP1 and STP2 redundant pathway (Right: Wildtype, Left: STP2 deletion mutant). The lack of STP2 is compensated by STP1, resulting in no change in the mean expression level of downstream genes (orange circles). However, expression noise that propagated from upstream changes differ due to changes in physical characteristics of upstream genes.

Notably, not all noise changes imply functional compensation, as noise change reflects changes in physical properties, such as upstream translational efficiency, and does not necessarily indicate functional compensation. If functional compensation occurs upstream, the genes must show noise change without mean change (noise-only change), due to changes in physical properties and compensation effects on the expression level. Therefore, we focused on genes that show noise-only change and investigated whether they are enriched in the common downstream genes of redundant gene pairs, such as STP1/2.

In this study, we aimed to estimate mean-change genes and noise-change genes in yeast using scRNA-seq data by comparing expression patterns in a single deletion mutant (ΔSTP1, ΔSTP2, ΔRTG1, ΔRTG3, ΔGLN3, ΔGAT1, ΔGZF3, and ΔDAL80; Table S1) with those in wildtype.

## Results and discussion

### Redundantly regulated target; STP1 and STP2

*STP1* and *STP2* are an ohnologous pair that arose from whole genome duplication, and thus share many downstream genes. Since it has been reported that STP1 and STP2 have redundant functions [7], those common targets are likely to be redundantly regulated. Regarding the downstream genes shared by STP1 and STP2, the proportion of genes that show mean expression change in the ΔSTP1 and ΔSTP2 was 36% and 25%, respectively. The remaining genes did not show mean expression change in the single deletion, which suggests functional compensation [4, 5, 6]. Based on our hypothesis, those redundantly regulated genes can be detected by noise-only change, even in the single deletion of *STP1* or *STP2*.

We examined whether mean change genes or noise-only change genes in ΔSTP1 and ΔSTP2 respectively were enriched in the common downstream genes of STP1 and STP2.

The mean change genes upon ΔSTP2 were not enriched in the downstream genes shared by STP1 and STP2 (*p* < 0.05, Fisher’s exact test; Fig. 2a). This result was consistent with a previous study showing that most downstream genes of STP2 were undetectable from single deletion of *STP2* (Reimand J et al., 2010). However, mean change genes upon ΔSTP1 were enriched in the downstream genes shared by STP1 and STP2 (*p* < 0.05, Fisher’s exact test; Fig. 2a). This result suggests that the deletion of *STP1* was not well compensated by STP2, although a previous study reported that deletion of *STP1* was compensated by STP2 [7]. Subsequently, we quantified noise considering heterogeneity due to cell cycle. Briefly, we clustered WT and mutant cells simultaneously by similar expression pattern and executed comparison within each clusters (Fig. S1 and see Material and methods). Because this analysis is vulnerable to the clustering methods, we keep this analysis as supplement. When we executed the same analysis with Fig. 2 but with considering cell cycle heterogeneity (Fig. S2a and see Materials and Methods), the same tendency with Fig 2a were maintained. The previously reported common interaction detected only by a double deletion was 64% in STP1 and 75% in STP2. The remaining genes were able to be detected even from a single deletion [4, 5, 6]. On the contrary, the noise-only change genes were enriched in the downstream genes shared by STP1 and STP2 in both ΔSTP1 and ΔSTP2 (*p* < 0.05, Fisher’s exact test; Fig. 3a). This result supports our hypothesis that the downstream genes of functionally redundant genes show noise-only change. When we eliminated cell cycle heterogeneity, the enrichment of noise-only change genes to common downstream genes of STP1 and STP2 disappeared in ΔSTP1, but a relatively high proportion of noise-only change genes (20%) were common downstream genes of STP1 and STP2 (Fig. S3a).

**Figure 2.**
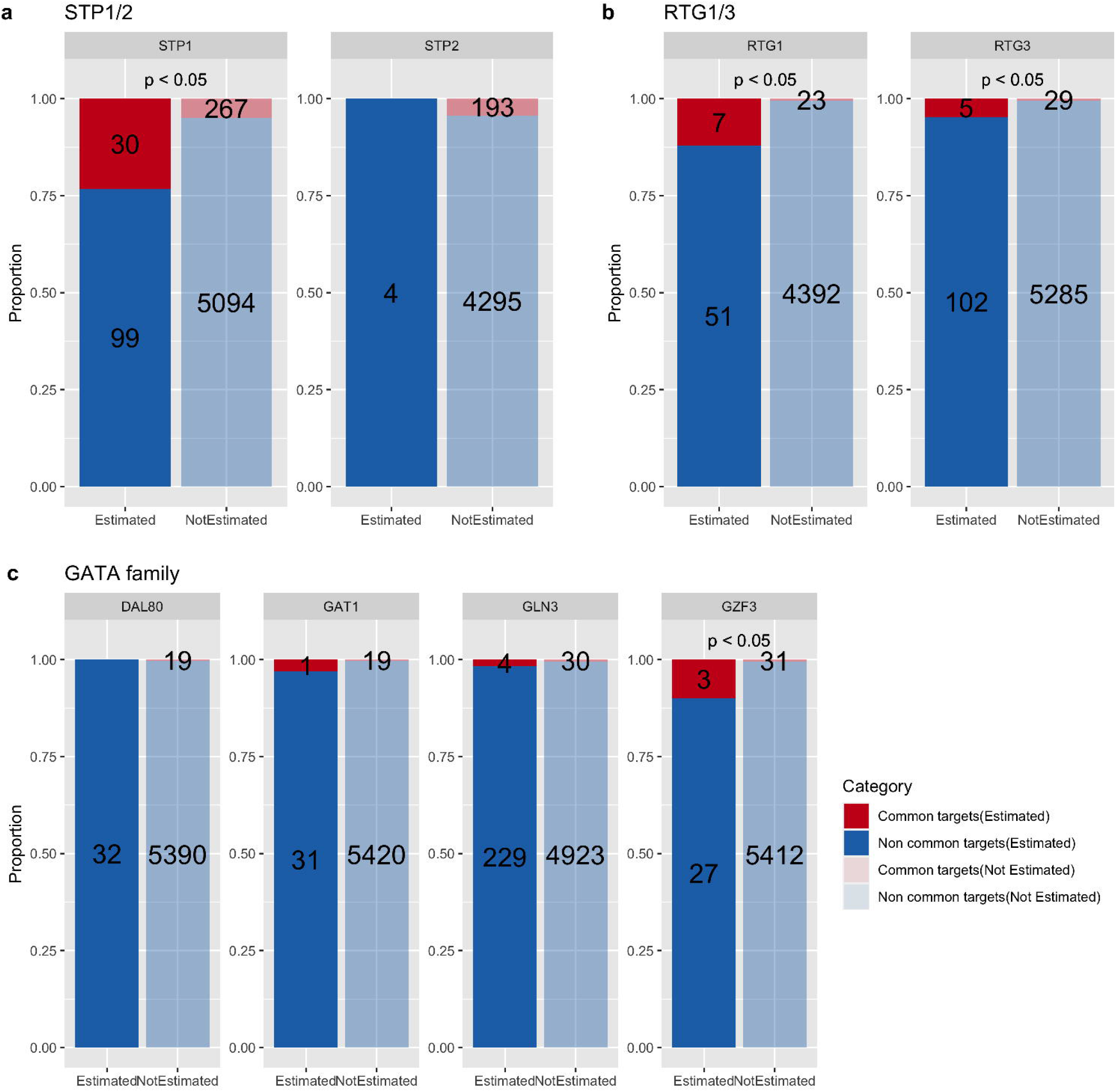
Enrichment test of mean change genes to known downstream genes shared by homologous groups. The bar graph shows whether the mean change genes in each deletion strain are enriched in the common downstream genes of the groups in which the deleted gene belongs. (a) Redundant gene group; STP1 and STP2. (b) Non redundant gene group; RTG1 and RTG3. (c) Possibly redundant paralogous group; GATA family (DAL80, GAT1, GLN3, and GZF3). The *p*-value above the bar graphs were calculated using Fisher’s exact test. The dark bars on the left show the proportion of known common interactions (red) and unknown common interactions (blue) in the mean change genes (FDR < 0.05). The light bars on the right show the proportion of known common interactions (red) and unknown common interactions (blue) in the non-mean change genes. The vertical axis represents the proportion, and the numbers on the bars show the number of genes in each category. If the red area on left is significantly larger than that on right, it indicates that mean change genes are enriched in common downstream genes.

**Figure 3.**
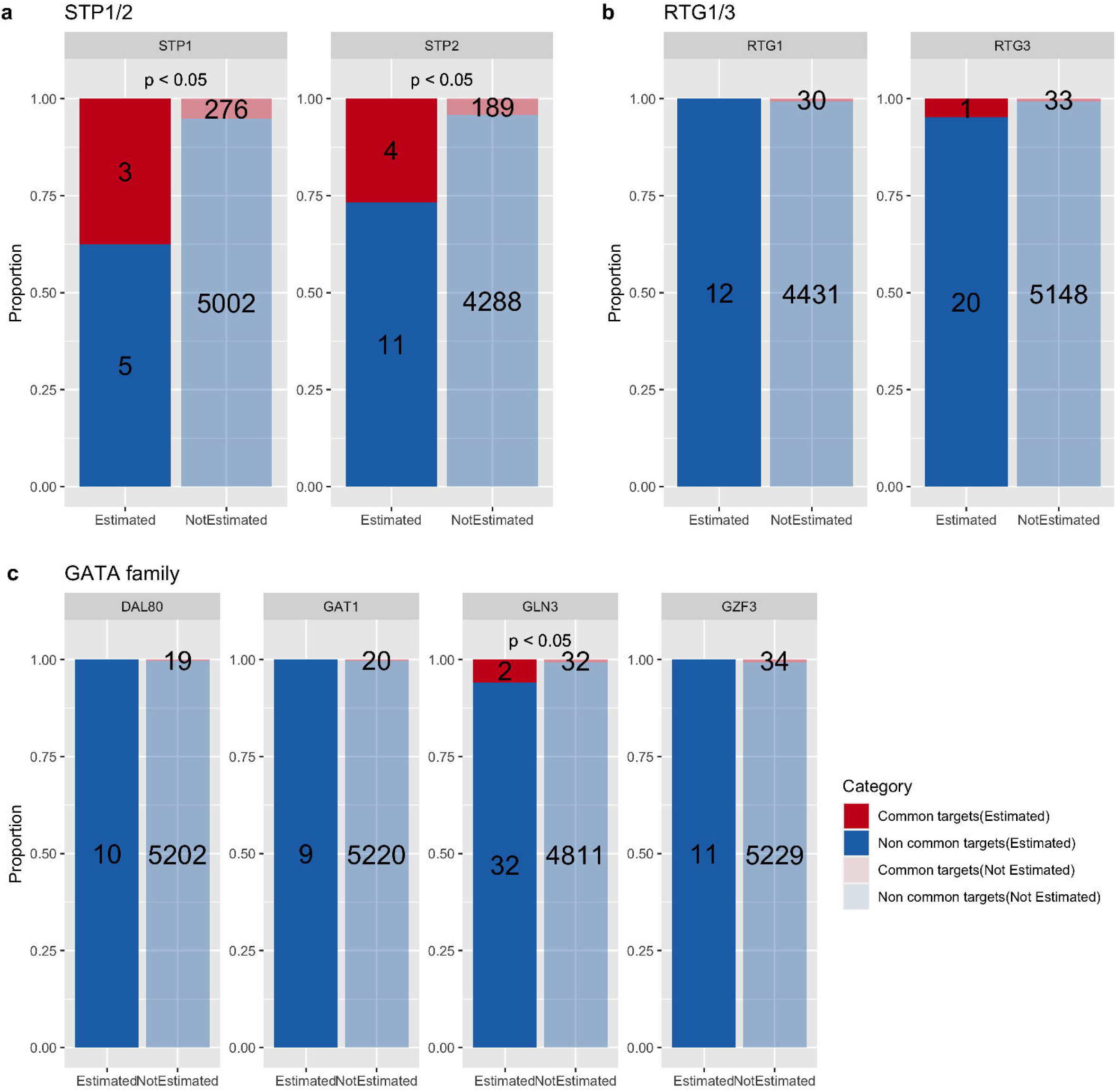
Enrichment test of noise-only change genes to known downstream genes shared by homologous groups. The bar graph shows whether the noise-only change genes in each deletion strains are enriched in the common downstream genes of the groups in which the deleted gene belongs. (a) Redundant gene group; STP1 and STP2. (b) Non redundant gene group; RTG1 and RTG3. (c) Possibly redundant paralogous group; GATA family (DAL80, GAT1, GLN3, and GZF3). The *p*-value above the bar graphs were calculated using Fisher’s exact test. The dark bars on the left indicate the proportion of known common interactions (red) and unknown common interactions (blue) in the noise-only change genes (FDR < 0.05). The light bars on the right indicate the proportion of known common interactions (red) and unknown common interactions (blue) in the non-noise-only change genes. The vertical axis represents the proportion, and the numbers on the bars show the number of genes in each category. If the red area on left is significantly larger than that on right, it indicates that noise-only change genes are enriched in common downstream genes.

### Non redundantly regulated target; RTG1 and RTG3

Next, as control, we investigated downstream genes shared by non-redundant genes. *RTG1* and *RTG3* are not a paralog pair, but they have significant overlap in downstream genes to form a complex [6]. We investigated whether mean change genes or noise-only change genes were enriched in the downstream genes shared by RTG1 and RTG3. The mean change genes were enriched in the common downstream genes in both ΔRTG1 and ΔRTG3 (*p* < 0.05, Fisher’s exact test; Fig. 2b). The results are reasonable because RTG1 and RTG3 do not have functional compensation and the perturbation of even one gene produces downstream effects. We observed the same trend even when we considered cell cycle heterogeneity (Fig. S2b). The noise-only change genes were not enriched in ΔRTG1 nor ΔRTG3 (*p* < 0.05, Fisher’s exact test; Fig. 3b). This is also consistent with the result considering cell cycle heterogeneity (Fig. S3b). These results suggest that RTG1 and RTG3 do not exhibit functional compensation.

### Possibly redundantly regulated target; GATA family

Next, we focused on the GATA family, *GAT1, GZF3, DAL80*, and *GLN3* [10], and investigated networks constituted from homologous gene groups. They are paralogs and possibly have redundant functions. We investigated whether these genes display functional compensation by estimating whether noise-only change genes are enriched in common downstream genes of the paralogous family. In ΔGZF3, the mean change genes were enriched in downstream genes shared by GZF3 and other GATA family members (*p* < 0.05, Fisher’s exact test; Fig. 2c). However, other deletion strains from the GATA family did not show enrichment of mean change genes compared to common downstream genes. This implies the functional compensation or merely non-activation of GATA family members under our study conditions. When we considered cell cycle heterogeneity, the genes did not show the enrichment (Fig. S2c). Thus, enrichment in ΔGZF3 in Figure 2c may indicate a false positive. The noise-only change genes were enriched in the common downstream genes of the GATA family only in ΔGLN3 (*p* < 0.05, Fisher’s exact test; Fig. 3c). Although a redundant pathway has not been reported between GLN3 and other genes, double deletion of *GLN3* and *GAT1* or *GLN3* and *DAL80* decreased growth rate [11, 12]. These results suggest the existence of functional compensation between GLN3 and GAT1 and/or GLN3 and DAL80. However, the noise change genes caused by ΔGAT1 or ΔDAL80 were not enriched in the common downstream genes. It has been reported that GATA family members were expressed under nitrogen limitation [10], thus the GATA family was necessarily required in nitrogen rich media, such as YPD, which was used in this study, resulting in no functional compensation. In addition, the small number of common downstream genes may also hinder target estimation by noise-only change. In Figure S3c, the strains did not show enrichment with common downstream genes with the GATA family when we considered cell cycle heterogeneity.

From the above results, we suggest that noise-only change captures redundantly regulated genes. Based on our concepts, the candidate of redundantly regulated genes, which was detected by noise-only change, must be common to the deletion strain of the redundant gene pair. Further, we detected three genes (*MUP1, CDC21, GAS3*) that show noise-only change in both ΔSTP1 and ΔSTP2, while two of these (*MUP1, CDC21*) are known as shared downstream genes of STP1 and STP2. We propose that *GAS3* is a new STP1/2 redundantly regulated candidate. Although an interaction between STP2 and *GAS3* has not been reported, an interaction between STP1 and *GAS3* was reported [6]. This supports that *GAS3* is a new redundantly regulated candidate by STP1/2. Notably, although the number of noise-only change genes common to ΔSTP1 and ΔSTP2 was three, the number of mean change genes common to ΔSTP1 and ΔSTP2 was zero. These results support our hypothesis that redundantly regulated genes that cannot be detected from mean change but are detected from noise-only change.

However, not all noise-only change genes are common to ΔSTP1 and ΔSTP2. The total number of noise-only change genes in ΔSTP1 is 15, while there are eight in ΔSTP2. As described above, the three genes overlap. Most noise-only change genes are detected only in ΔSTP1 or ΔSTP2. We speculate that this asymmetric noise-only change is caused by (1) artifact of FDR threshold, (2) asymmetric compensation, or (3) compensation by other genes. First, since the extent of noise difference caused by gene deletion is attributed to physical properties of the deleted gene, such as translational efficiency [9], the threshold must differ gene by gene. Applying the same cut off to all genes leads to increased false negative results. The precise modeling of noise propagation must improve detection of redundant regulation. Second, asymmetric functional compensation might be considered. For example, the deletion of *STP2* is compensated by STP1, but the deletion of *STP1* is not compensated by STP2 with fewer interactions. In such situations, the gene shows mean change upon ΔSTP1 and on the contrary the gene shows noise-only change upon ΔSTP2. Third, functional compensation by other genes might be considered. We assumed that functional compensation is likely to occur between paralogous pairs due to their functional similarity, but there are many synthetic effects between *STP1/2* and other genes [11]. However, a synthetic effect does not necessarily translate to functional compensation, but those interactions suggest that a lack of *STP1* or *STP2* functionality may possibly be compensated by the other genes.

### Novel downstream candidates of STP1 and STP2 detected from noise change

We investigated noise change genes, including mean change, to determine whether noise difference tests narrow down the reasonable downstream genes. We hypothesized that noise-only change genes shared by the deletion strain of a paralogous pair, such as ΔSTP1 and ΔSTP2, are likely to be the target of functional compensation. Other noise change genes can be recognized as downstream candidates. Interestingly, both known STP2 downstream genes and novel STP2 downstream candidates detected from noise change were enriched in transport-related genes (*p* < 0.05; SGD GO Term Finder; Process; Table 1). Therefore, it can be argued that the noise difference test has targeted reasonable candidates. *ARG1*, a novel downstream candidate of STP2, was confirmed as a downstream STP1 gene [6], which implies redundant regulation by STP1 and STP2.

**Table 1.**
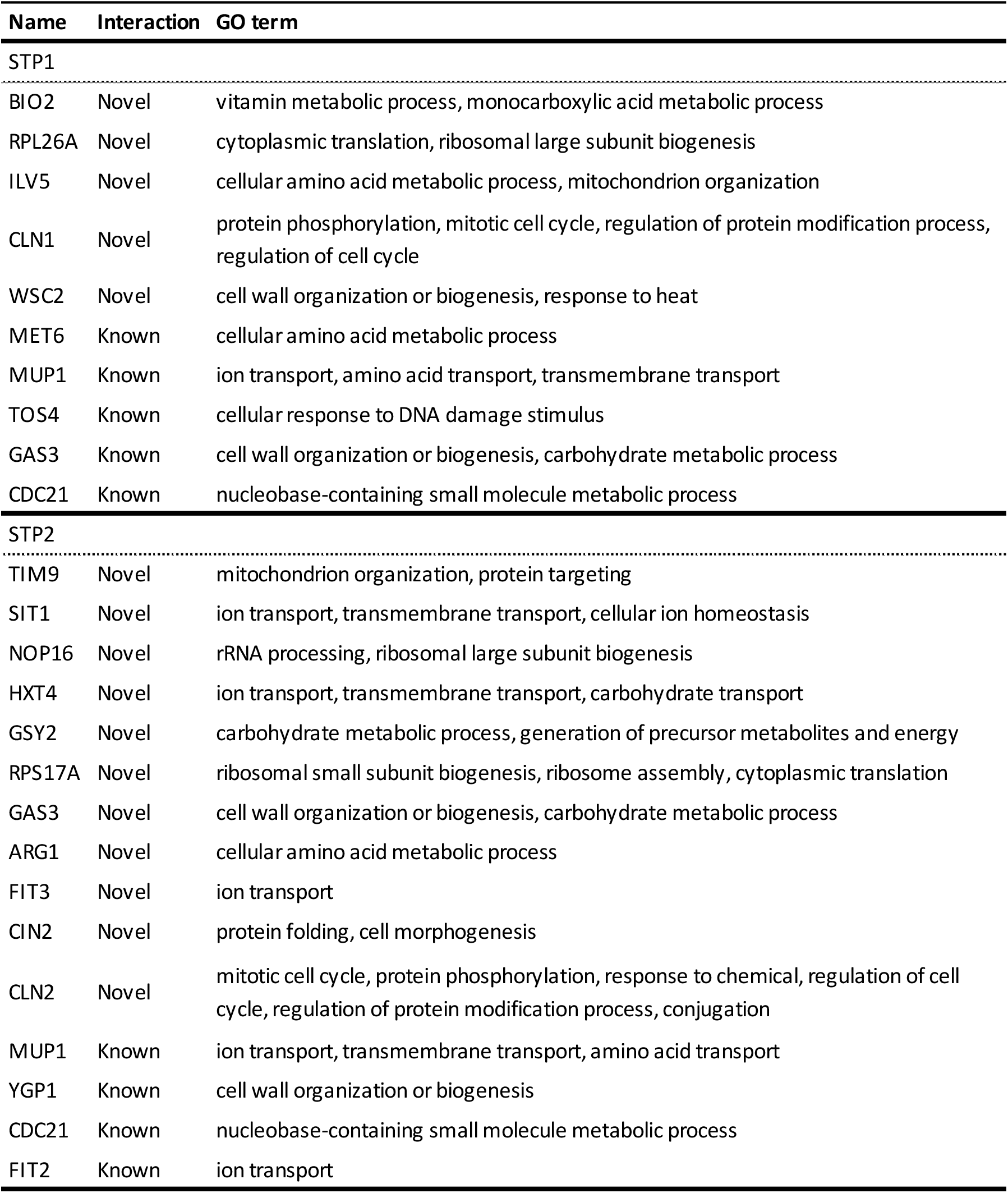
Noise-change genes in ΔSTP1 and ΔSTP2. Name: Genes that showed noise change in ΔSTP1 or ΔSTP2 are listed; Interaction: Noise change genes that had not been reported as STP1 or STP2 downstream in Yeastract are labeled as “Novel”; GO term: GO terms retrieved from SGD GO slim mapper (Yeast GO-Slim process). In the novel STP1 candidates, there were no enriched GO terms (p < 0.01; SGD GO term finder process). In the novel STP2 candidates, GO terms were enriched in transport-related genes, in which STP2 downstream genes are involved (p < 0.01; SGD GO Term Finder; Process).

In this study, we executed two types of analysis. (1) The conservative difference test, which quantified values from all cells (Fig. 2 and 3), and (2) the progressive difference test, which quantified values from cells belonging to the same cluster (Fig. S2 and S3). In the later progressive analysis, as cell cycle heterogeneity is excluded from noise quantification, we can quantify that the noise change originated from a molecular process (but the results are affected by clustering). In the former conservative analysis, *CLN1*, which is related to cell cycle regulation and does not interact with STP1, showed noise change upon ΔSTP1, but in the later progressive analysis, CLN1 did not show noise change upon ΔSTP1 (Table S2). In the same way, *CLN2*, which is related to the cell cycle and does not interact with STP2, showed noise change upon ΔSTP2, but when we consider cell cycle heterogeneity, CLN2 did not show significant noise change (Table S2). This implies that the noise change of CLN1 and CLN2 originated from differences in cell cycle heterogeneity that are out of our interests. We consider that the consistent results of conservative and progressive analysis should be reliable. We found four consistent genes in ΔSTP2 that show noise change, both from conservative and progressive analysis (*CDC21, SIT1, FIT2, FIT3*). *FIT2* and *CDC21* are known as shared downstream genes of STP1 and STP2. *SIT1* and *FIT3* are possibly newly downstream candidates of STP2, and work as transporters, similarly to many known downstream genes of STP2. In addition, *SIT1* and *FIT3* are known as downstream genes of STP1, and the interaction between STP2 and *SIT1* or *FIT3* could not be detected from mean difference, although detected from noise change. Thus, we suggest they are redundantly regulated by STP1 and STP2. In summary, we detected four novel gene candidates that are redundantly regulated by STP1 and STP2 (*ARG1, FIT3, SIT1, GAS3*), even from the single deletion strain of STP2. Three of these (*FIT3, SIT1, GAS3*) are constant, as we observed consistent results across progressive and conservative analyses. Interestingly, ARG1 is involved in the arginine biosynthesis process [13], and it was previously unknown that STP2 affects this process. Thus, our results suggest that STP2 may be involved in the arginine biosynthesis process.

Based on the above results, we constructed the gene interaction network shared by STP1 and STP2 (Fig. 4). *MUP1, FIT2*, and *YGP1* were only detected from mean change in the double deletion of *STP1* and *STP2* [4, 5]. Namely, they cannot be detected in single deletions of *STP1* or *STP2*. However, in this study, those hidden interactions could be detected by the noise-only change, even from the single deletion of *STP2*. In addition, several new STP2 downstream genes were detected by noise change. We found that some of these were also downstream genes of STP1, which supports that they are redundantly regulated by STP1/2 (genes indicated by blue and red arrow in Fig. 4). Here, we discuss about the reason why those novel STP2 downstream genes that have redundant interaction with STP1, had not been found by ΔSTP1ΔSTP2 nor ΔSTP2. If the effect of STP1 is much larger than that of STP2 in redundant pathway, the mean expression change of downstream genes estimated in ΔSTP1ΔSTP2 may not differ from that in ΔSTP1 due to the subtle effect of STP2. Namely, the detection of asymmetric functional compensation is difficult, even from the double deletion, unless we use noise-only change. Furthermore, we found that several newly estimated genes that are redundantly regulated by STP1/2 have positive or negative synthetic interaction (see methods) or similar interaction patterns (profile similarity) with known redundantly regulated genes. This also validates the novel interactions.

**Figure 4.**
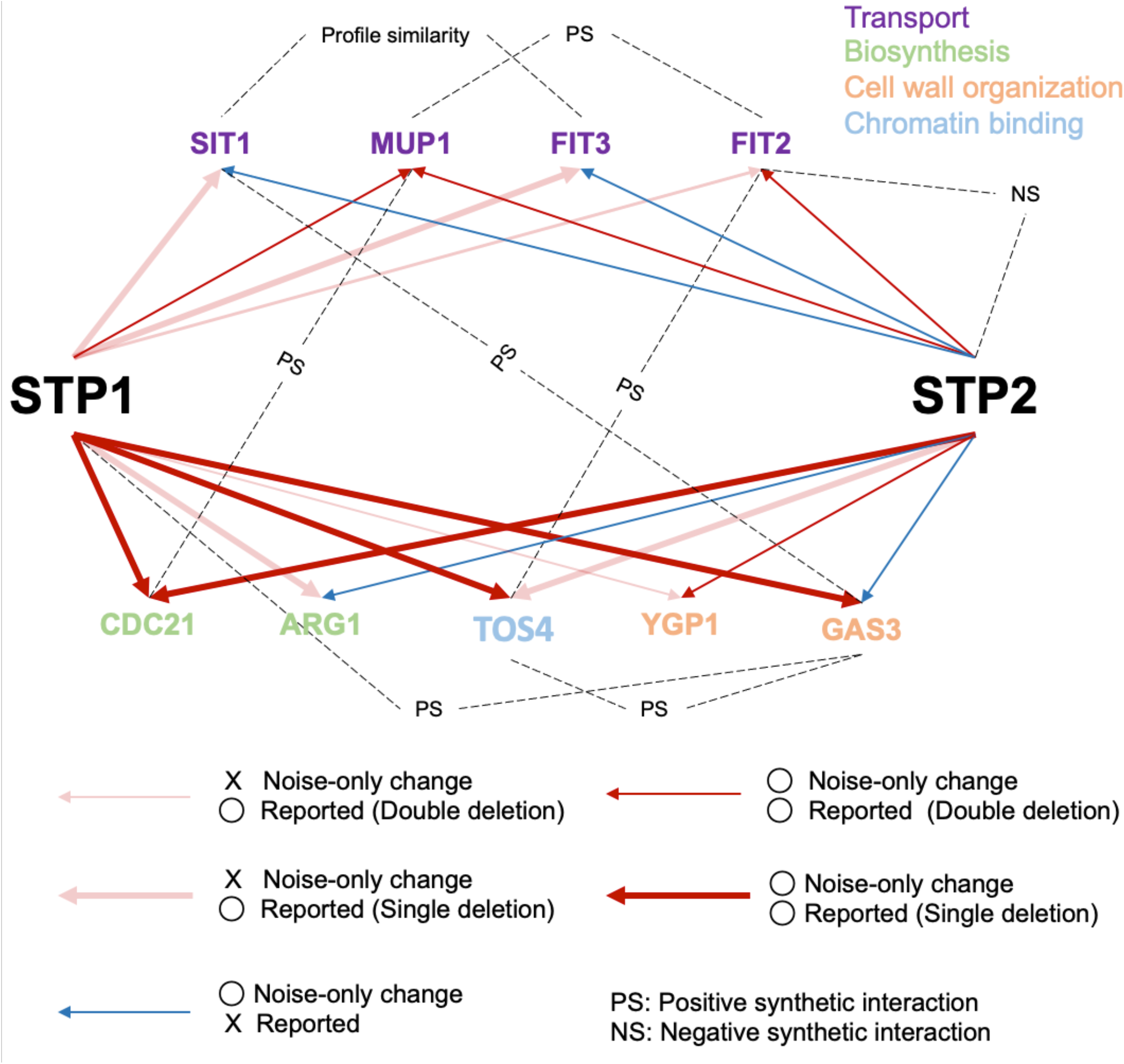
Gene regulatory network shared by STP1 and STP2. Dark arrows suggest the interaction detected from noise-only change. Red and blue arrows imply known and novel interactions, respectively. Light red arrows suggest known interactions that cannot be detected from noise-only changes in this study. Wide red arrows imply that the interaction was previously detected from a single deletion, and the thin red arrows imply the interaction was previously detected from a double deletion. Dashed lines indicate a synthetic effect, which are negative synthetic interaction (NS) or positive synthetic interaction (PS). The color of gene names implies GO terms retrieved from SGD GO Slim mapper. SIT1 and FIT3 depicts profile similarity defined by the similarity of synthetic interactions.

Conventionally, redundantly regulated genes are detected by assessing mean expression change in the double deletion of redundant genes, in order to avoid functional compensation. From this analysis, we propose that redundantly regulated genes can be detected by assessing noise-only change, even in the single deletion of one of the redundant genes. Our new approach will provide a novel framework for revealing complex gene networks. As a limitation of this study, because it is necessary to quantify the variation of expression levels in homogeneous cells, as we showed in this study (Fig. S1 and see Materials and Methods), its application is limited to unicellular organisms at this stage. Furthermore, the quantified noise includes intrinsic noise and extrinsic noise, which later one is not our interests. More accurate interaction analysis should be possible by eliminating extrinsic noise and focusing on intrinsic noise. In this study, we roughly excluded extrinsic noise by clustering scRNA-seq data (Fig. S1). We propose that exact modeling of extrinsic and intrinsic noise using scRNA-seq data will allow for more accurate noise difference estimation. We consider that noise difference tests can be applied as a new strategy for data-driven network estimation that can capture even functional compensation.

## Materials and methods

### Expression data

We used single-cell RNA-seq data in yeast, including eight different TF deletion strains (ΔSTP1, ΔSTP2, ΔRTG1, ΔRTG3, ΔGLN3, ΔGAT1, ΔGZF3, and ΔDAL80) and wild type (WT) as a control [14]. ΔSTP1 and ΔSTP2 were used to investigate a redundant gene pair. ΔRTG1 and ΔRTG3 were used to investigate a non-redundant gene pair that shares many target genes. ΔGAT1, ΔGZF3, ΔDAL80, and ΔGLN3 were used to investigate paralogues gene groups, from which redundant functions have not been reported. Each strain had six biological replicates, which were independently constructed and cultured under YPD conditions. Six of the deleted TFs were paralogs (BLASTP search, E-value < 10^−10^) and two of these were singletons (Table S1). We obtained transcript counts from individual cells, which passed quality control, followed by removal of doublets according to Jackson *et al*., 2019 [14].

### Elimination of cell cycle heterogeneity

In order to precisely quantify the changes in expression noise caused by gene deletion, cell heterogeneity due to cell cycle should be eliminated. Therefore, we divided cells into groups exhibiting similar expression patterns. Two comparative strains were clustered simultaneously. This enabled us to compare two strains within a cluster. In order to cluster cells by their expression state, first, we normalized cell specific biases using scran normalization methods [15]. Subsequently, we constructed shared nearest neighbor graphs [16] from normalized counts, and clustered these using Louvain methods [17]. As a result, the strains were grouped into five or six clusters (Fig. S1). Although cells of two strains were randomly distributed in the clusters, expression patterns of *DSE2*[18], *PIR1* [19], and *HTB* [20], known as cell cycle marker genes, were biased by clusters (Fig. S1). Thus, cells were thought to be clustered by cell cycle. Since each cluster represents a specific cell state, two strain comparisons within a cluster was not thought to be affected by heterogeneity due to cell cycle. Since the comparison was executed by cluster, noise (mean) difference results for one gene in a strain equated to the number of clusters. If a gene showed significant change (FDR < 0.01; Bonferroni correction) in at least one cluster, we considered the gene as a downstream gene candidate of the deleted gene. The results were shown in Fig. S2 - S7. Since these methods require large sample sizes, clustering was not executed in a single deletion mutant GLN3 with a small sample size. Detected genes in this method are not robust as they are affected by clustering.

### Filtration of technical noise

As we described above, each strain includes six biological replicates constructed from different experiments. These replicates were utilized to estimate technical noise under the assumption in which replicate specific variation is technical artifact, and common variation between replicates is biological noise. This filtration was executed using the hierarchical bayesian model in BASiCS R package [21, 22].

### Expression level comparison

To test the hypothesis that functional compensation is not detected by mean change but can be detected by noise change, we compared differences in mean expression levels for each gene between wildtype and deletion mutant with differences in expression noise. The mean expression level and expression noise were estimated using the hierarchical Bayesian model in BASiCS R package [21, 22]. All parameters of MCMC were set to the recommended values in the BASiCS R package manual. Since the expression noise used in the comparison was a residual measure of variability that is not correlated with mean expression level, expression noise change is independent from mean change. The details were reported in Eling *et al* 2018 [21](residual over-dispersion). After comparing the mean with noise respectively, genes showing a significant change with EFDR less than 0.01 were estimated as downstream genes of the deleted gene.

### Validation of estimation results

To investigate whether estimated downstream genes were enriched in reported downstream genes, we used the experimentally confirmed interaction database, Yeastract [23]. We retrieved the regulation matrix from Yeastract. Only expression evidence detected under unstressed log-phase growth conditions were included.

### Positive/Negative synthetic interaction and profile similarity

As a result of this study, we constructed shared genetic interaction networks of STP1 and STP2. To support the validity of our results, we used other interaction data, positive or negative synthetic interaction and profile similarity reported by Costanzo and colleagues [11]. The positive synthetic interaction implies that the double deletion mutant is less severe than expected from each single deletion mutant. The negative synthetic interaction implies that the double deletion mutant is much more severe than expected from each single deletion mutant. The profile similarity implies the similarity of synthetic effects measured by Pearson correlation [11]. The gene pair that exhibited a similarity score greater than 0.2 was recognized as “Profile similarity” in our study.

## Abbreviations

GRN: Gene regulatory network
TF: Transcription factor

## Acknowledgements

We would like to thank Hisao Moriya for valuable comments on our study and all the members of the Makino Laboratory for helpful discussions.

## Data availability

All data analyzed during this study are included in this manuscript.

Scripts used in this analysis have been deposited in https://zenodo.org/record/5041324#.YNu_yBMzbb8

## Author contribution

T.I. and T.M. conceived and designed the project. T.I. conducted experiments. T.I. and T.M. analyzed the data. T.I. and T.M. wrote the paper.

