## Supplement file for "Capturing hidden regulation based on noise change of gene expression level from single cell RNA-seq in yeast"

**Table S1. Eight deletion strains and associated paralogs.**

| Strain | Paralog |
|---|---|
| ΔSTP1 | STP2 |
| ΔSTP2 | STP1 |
| ΔDAL80 | GLN3, GAT1, GZF3 |
| ΔGLN3 | DAL80, GAT1, GZF3 |
| ΔGAT1 | DAL80, GLN3, GZF3 |
| ΔGZF3 | DAL80, GLN3, GAT1 |
| ΔRTG1 | Nothing |
| ΔRTG3 | Nothing |

**Table 2. Noise change genes in ΔSTP1 and ΔSTP2 considering cell cycle heterogeneity.** Name: Genes that showed noise change considering cell cycle heterogeneity are listed; Interaction: Noise change genes that had not been reported as STP1 or STP2 downstream in Yeastract are labeled as "Novel"; GO terms: GO term retrieved from SGD GO slim mapper (Yeast GO-Slim process). In novel STP1 candidates, there were no enriched GO terms ($p < 0.01$; SGD GO term finder process). In novel STP2 candidates, GO terms were enriched in transport-related genes, in which STP2 downstream genes are involved ($p < 0.01$; SGD GO Term Finder; Process).

| Name | Interaction | GO term |
|---|---|---|
| STP1 | | |
| BIO2 | Novel | vitamin metabolic process, monocarboxylic acid metabolic process |
| RPL26A | Novel | cytoplasmic translation, ribosomal large subunit biogenesis |
| ILV5 | Novel | cellular amino acid metabolic process, mitochondrion organization |
| CLN1 | Novel | protein phosphorylation, mitotic cell cycle, regulation of protein modification process, regulation of cell cycle |
| WSC2 | Novel | cell wall organization or biogenesis, response to heat |
| MET6 | Known | cellular amino acid metabolic process |
| MUP1 | Known | ion transport, amino acid transport, transmembrane transport |
| TOS4 | Known | cellular response to DNA damage stimulus |
| GAS3 | Known | cell wall organization or biogenesis, carbohydrate metabolic process |
| CDC21 | Known | nucleobase-containing small molecule metabolic process |
| STP2 | | |
| TIM9 | Novel | mitochondrion organization, protein targeting |
| SIT1 | Novel | ion transport, transmembrane transport, cellular ion homeostasis |
| NOP16 | Novel | rRNA processing, ribosomal large subunit biogenesis |
| HXT4 | Novel | ion transport, transmembrane transport, carbohydrate transport |
| GSY2 | Novel | carbohydrate metabolic process, generation of precursor metabolites and energy |
| RPS17A | Novel | ribosomal small subunit biogenesis, ribosome assembly, cytoplasmic translation |
| GAS3 | Novel | cell wall organization or biogenesis, carbohydrate metabolic process |
| ARG1 | Novel | cellular amino acid metabolic process |
| FIT3 | Novel | ion transport |
| CIN2 | Novel | protein folding, cell morphogenesis |
| CLN2 | Novel | mitotic cell cycle, protein phosphorylation, response to chemical, regulation of cell cycle, regulation of protein modification process, conjugation |
| MUP1 | Known | ion transport, transmembrane transport, amino acid transport |
| YGP1 | Known | cell wall organization or biogenesis |
| CDC21 | Known | nucleobase-containing small molecule metabolic process |
| FIT2 | Known | ion transport |

**Figure S1. Cells clustered by cell cycle state.** Dimension composition was conducted using UMAP. First row: Clustering results; Second row: genotype mapping. Yellow and green plots represent mutant deletion and wildtype, respectively; Third row: *PIR1* expression pattern; Fourth row: *DSE2* expression pattern; Fifth row: *HTB* expression pattern.

**Figure S2. Enrichment test of mean change gene with elimination of cell cycle effect to known downstream genes shared by homologous groups.** In genes showing significant mean change (FDR < 0.01; Bonferroni correction), at least one cluster was labeled as "Estimated". The representation of this figure is the same as in Figure 2.

**Figure S3. Enrichment test of noise change gene with elimination of cell cycle effect to known downstream genes shared by homologous groups.** In genes showing noise-only change (FDR < 0.01; Bonferroni correction), at least one cluster was labeled as "Estimated". The representation of this figure is the same as in Figure 3.

### Figure S1

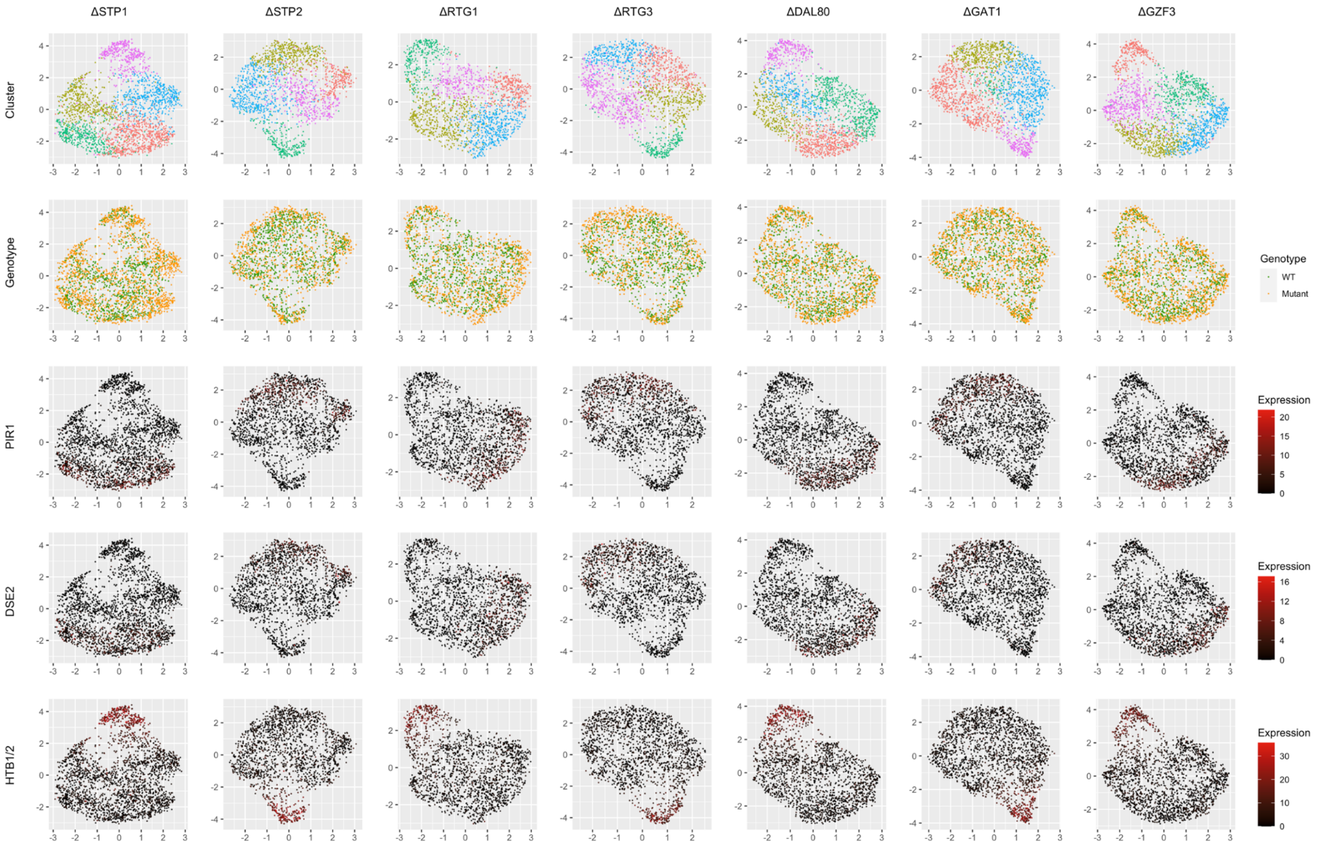

ΔSTP1
ΔSTP2
ΔRTG1
ΔRTG3
ΔDAL80
ΔGAT1
ΔGZF3
Cluster
Genotype
PIR1
DSE2
HTB1/2
Genotype
WT
Mutant
Expression
Expression
Expression

Figure S2

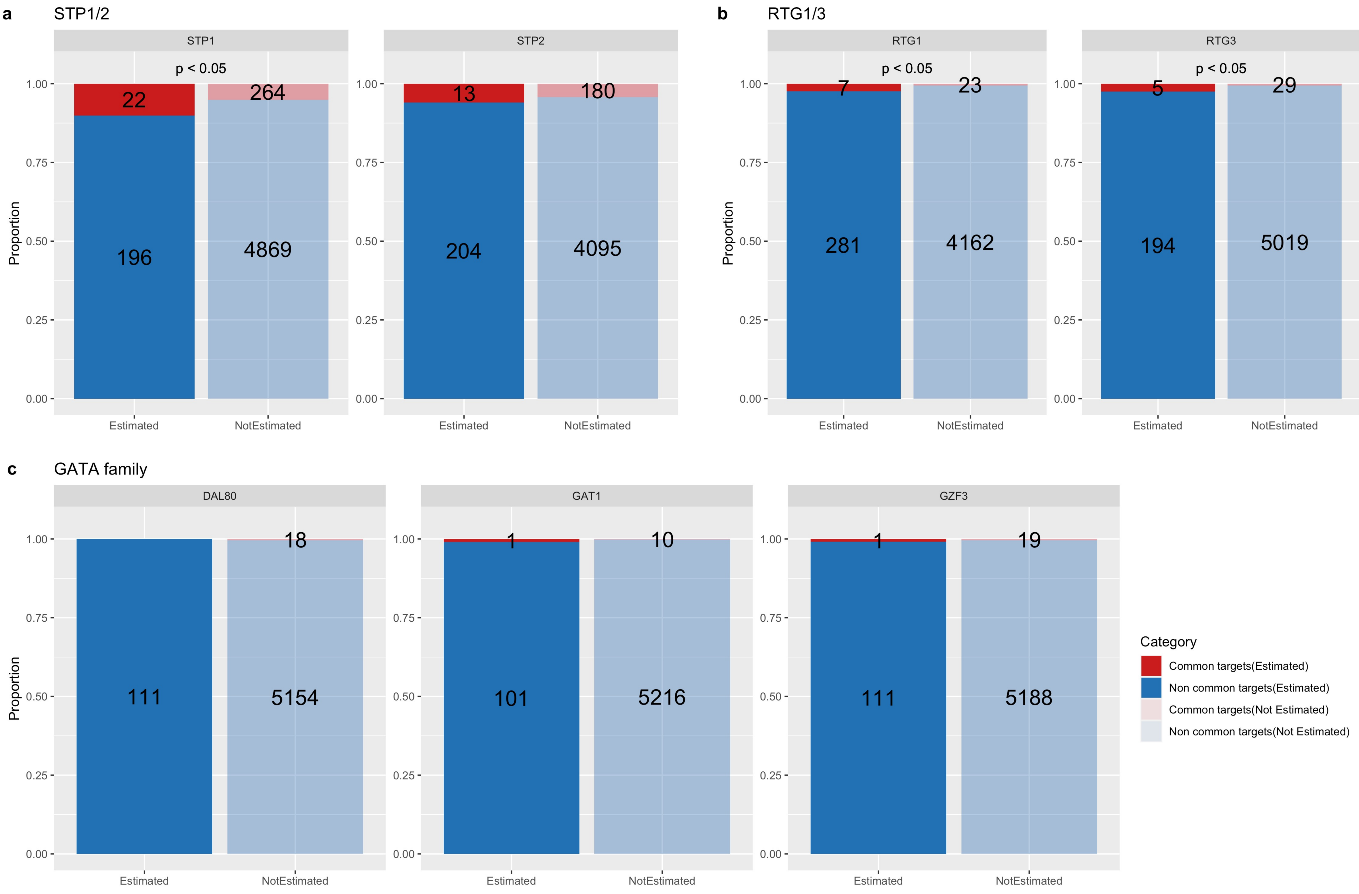
a
STP1/2
STP1
STP2
p < 0.05
22
264
196
4869
13
180
204
4095
Proportion
1.00
0.75
0.50
0.25
0.00
Estimated
NotEstimated
b
RTG1/3
RTG1
RTG3
p < 0.05
7
23
281
4162
5
29
194
5019
c
GATA family
DAL80
GAT1
GZF3
18
111
5154
1
10
101
5216
1
19
111
5188
Category
Common targets(Estimated)
Non common targets(Estimated)
Common targets(Not Estimated)
Non common targets(Not Estimated)

### Figure S3

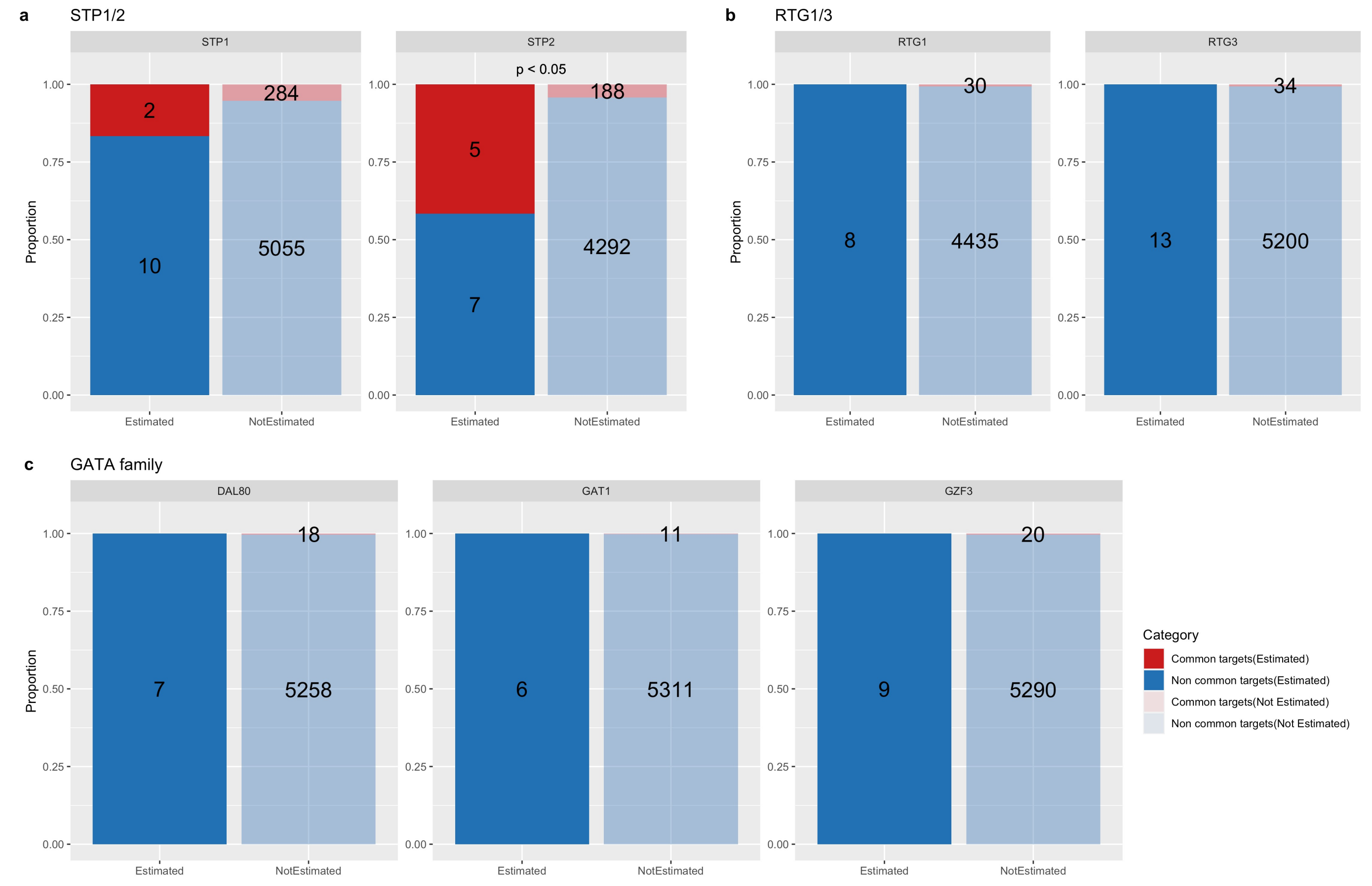

a
STP1/2
STP1
STP2
p < 0.05
Proportion
1.00
0.75
0.50
0.25
0.00
2
10
284
5055
5
7
188
4292
Estimated
NotEstimated
b
RTG1/3
RTG1
RTG3
8
30
4435
13
34
5200
c
GATA family
DAL80
GAT1
GZF3
7
18
5258
6
11
5311
9
20
5290
Category
Common targets(Estimated)
Non common targets(Estimated)
Common targets(Not Estimated)
Non common targets(Not Estimated)
